# The circadian clock and darkness control natural competence in cyanobacteria

**DOI:** 10.1101/822627

**Authors:** Arnaud Taton, Christian Erikson, Yiling Yang, Benjamin E. Rubin, Scott A. Rifkin, James W. Golden, Susan S. Golden

## Abstract

Natural genetic competence-based transformation contributed to the evolution of prokaryotes, including the cyanobacterial phylum that established oxygenic photosynthesis. The cyanobacterium *Synechococcus elongatus* is noted both as a model system for analyzing a prokaryotic circadian clock and for its facile, but poorly understood, natural competence. Here a genome-wide screen aimed at determining the genetic basis of competence in cyanobacteria identified all genes required for natural transformation in *S. elongatus*, including conserved Type IV pilus, competence-associated, and newly described genes, and revealed that the circadian clock controls the process. The findings uncover a daily program that determines the state of competence in *S. elongatus* and adapts to seasonal changes of day-length. Pilus biogenesis occurs daily in the morning, but competence is maximal upon the coincidence of circadian dusk and the onset of darkness. As in heterotrophic bacteria, where natural competence is conditionally regulated by nutritional or other stress, cyanobacterial competence is conditional and is tied to the daily cycle set by the cell’s most critical nutritional source, the Sun.

---

Horizontal exchange of genetic material through conjugation, phage transduction, and natural competence shaped the evolution of the prokaryotic domain of life^1^. The phenomenon of natural competence was famously discovered by Griffith in 1928 in the Gram-positive bacterium *Streptococcus pneumoniae*^2^. In Gram-negative bacteria such as *Vibrio cholerae* the core apparatus of DNA uptake is the type IVa pilus machine (T4PM) and an assemblage of competence-associated proteins^3^. The T4PM is composed of an extracellular pilus fiber (T4P) and a cell envelope–spanning complex called the basal body^4^. The major pilin PilA, processed by the prepilin peptidase PilD, makes a pilus that extends and retracts through the addition and removal of pilin subunits at the pilus base, powered by the assembly and disassembly adenosine triphosphatases PilB and PilT, respectively. The exogenous DNA (eDNA) that enters the periplasm through the T4PM is pulled in by ComEA, a DNA binding protein, and crosses the inner membrane through the ComEC channel. Once the eDNA, processed to single stranded DNA (ssDNA), reaches the cytoplasm, Ssb and DprA bind to the ssDNA and protect it from degradation; DprA also recruits RecA, which recombines the ssDNA with the chromosome. Although common mechanisms of DNA uptake and processing are thought to be shared by most transformable bacteria, numerous questions remain unanswered and the regulatory mechanisms that control competence vary among transformable species^5,6^.

*Synechococcus elongatus* PCC 7942 is a model cyanobacterium for the study of circadian rhythms^7^ and a platform strain for the production of biochemicals^8^. *S. elongatus* has been known to be naturally transformable since 1970^9^, but the mechanisms by which it or any other cyanobacterium takes up DNA from the environment are still poorly understood^10-12^. Cyanobacteria constitute a diverse phylum of prokaryotes that established oxygenic photosynthesis at least 3 billion years ago^13^ and contribute a large fraction of primary production in the oceans^14^. In contrast to other bacteria, cyanobacteria rely on light as a primary source of energy; accordingly, their cellular activities respond strongly to the presence and absence of light^15^. To synchronize their activities over a diel (24-h day-night) cycle, cyanobacteria use an endogenous circadian clock^7^. The proteins KaiA, KaiB, and KaiC comprise the core oscillator of the clock, which programs a ∼24 h cycle marked by the phosphorylation state of KaiC, whose autokinase and phosphatase activities are modulated by KaiA and KaiB. To synchronize the oscillator with the Earth’s diel cycle, input pathways that involve KaiA, KaiC, and the histidine kinase CikA monitor the light availability through cellular redox states and energy status^16,17^. To regulate circadian processes, an output pathway relays KaiC phosphorylation states to a two-component system with 2 antagonistic histidine kinases, CikA and SasA, and the response regulator RpaA^18^. RpaA serves as a master transcription factor that binds more than 134 transcript targets^19^ and enables complex circadian gene expression patterns by regulating a cascade of interdependent sigma factors^20^.

The cyanobacterial clock is known to regulate global gene expression^21^, chromosome compaction^22^, cell division^23,24^, and metabolite partitioning^15^. Here we screened a dense library of *S. elongatus* barcoded transposon mutants (RB-TnSeq)^25^ to uncover the genetic basis and regulation of natural competence in *S. elongatus*. Our data show that competence is controlled by the circadian clock and follows a model of external coincidence, wherein the external stimulus, darkness, must fall at the time of the dusk circadian peak to maximize competence, and that the regulation of this process adapts to the seasonal change of day-length.

## The Machinery

In order to globally identify all genes that enable natural competence, we transformed an *S. elongatus* RB-TnSeq library^25,26^ with eDNA carrying selectable markers that recombine into the *S. elongatus* chromosome at neutral sites (NS)^27^. We reasoned that transposon mutants that are incompetent for transformation would be selectively missing from the transformed population. Mutant colonies were collected as pools from selective and non-selective control conditions and the barcodes were sequenced. The relative abundance of mutants for each represented gene in both conditions was interpreted as a measure of fitness with a corresponding level of confidence (T-value) for each locus^26^ (Fig. 1a, Supplementary Table 1). Several functional categories that negatively or positively affect natural competence were represented. Genes whose loss significantly improves transformation encode homologs of Argonaute and Cas4. Strikingly, nearly all genes that encode the T4PM are necessary for cells to take up DNA (Fig. 1a,b), experimentally revealing that, as for other phyla of bacteria, natural competence in *S. elongatus* is mediated by the T4PM and specialized competence proteins. Competence-specific factors include ComEA, ComEC, ComF, and DprA, which are implicated in binding, pulling, and processing exogenous dsDNA as ssDNA in the periplasm, and translocating it to the cytoplasm. The RB-TnSeq data were confirmed by transformation assays performed on selected knockout mutants (Fig. 1c).

**Fig. 1.**
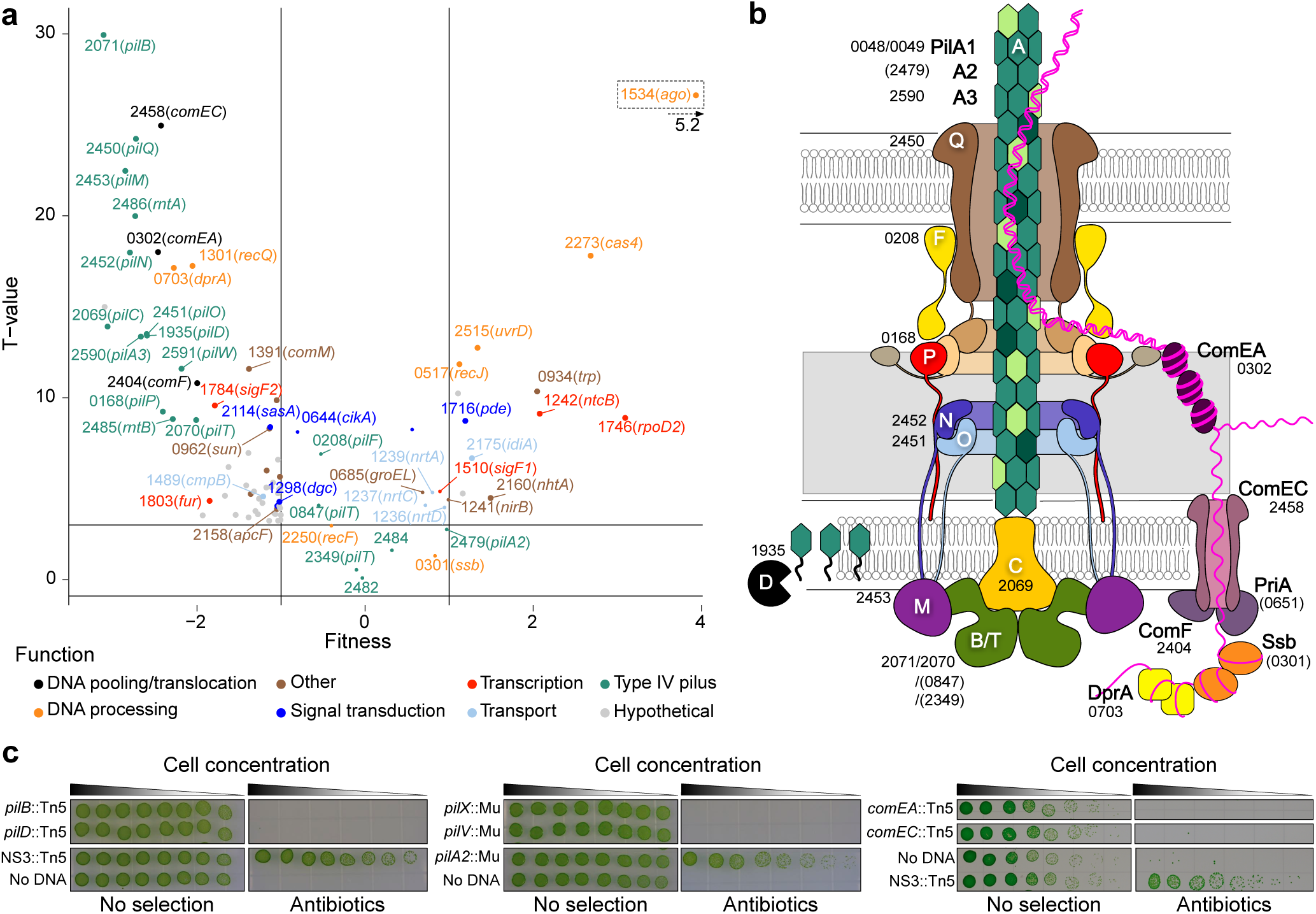
T4PM and competence proteins comprise the machinery for natural competence in cyanobacteria. **a**, Fitness values and confidence levels associated with barcoded transposon mutants for natural transformation. Loci that are essential for or strongly contribute to natural competence are characterized by a low fitness with a high T-value, whereas improved transformation following knockout of a locus generates a high fitness with a high T-value. Fitness values were calculated for 1885 loci, from 3 independent experiments, representing a total of 82,495 distinct mutants obtained under selective conditions and 90,872 under control conditions. With the exception of a few selected loci representing selected functional categories, only those whose loss of function had a strong and significant fitness effect are displayed on this plot. The data points were labeled with *S. elongatus* locus tag (Synpcc7942_) numbers and gene names in parentheses. **b**, Model of the T4PM based on a depiction by Chang et al^4^ in which competence-associated proteins are labeled with *S. elongatus* locus tag numbers. T4PM or competence protein homologs that are not required for natural competence are in parentheses. **c**, Transformation assays performed on insertion knockout mutants of selected loci. Strains of *S. elongatus* that carry a chromosomal insertion in loci known to not affect natural competence, *pilA2* or neutral site 3 (NS3), served as positive controls for transformation, and those same strains to which no eDNA was added served as negative controls.

Most of the identified proteins are conserved among cyanobacteria (Fig. 2a), even though only a few species are known to be naturally competent. Several novel genes are required for competence in *S. elongatus* but not widely conserved among cyanobacteria. These include two pairs of genes, *pilA3* and Synpcc7942_2591, and Synpcc7942_2486 and Synpcc7942_2485, whose products carry a type IV pilin-like signal peptide (Fig. 2a,b). While neither PilA3 nor Synpcc7942_2591 is encoded in other naturally competent cyanobacterial species, transformation assays demonstrated that both genes are required for competence in *S. elongatus* (Fig. 2c). In addition, PilA3 overexpression leads to increased transformation efficiency (Extended Data Fig. 1). Sequence similarities, a characteristic prokaryotic N-terminal methylation motif, and structural predictions clearly identified PilA3 as a pilin subunit. The Synpcc7942_2591 protein has no relatives with a known function but carries a PilW T4P assembly domain and so we designated it PilW. The roles of Synpcc7942_2486 and Synpcc7942_2485 are enigmatic; orthologs for Synpcc7942_2486 were found in competent strains but orthologs for Synpcc7942_2485 were not. Transformation assays showed that each gene is **r**equired for **n**atural **t**ransformation of *S. elongatus* and we named them *rntA* and *rntB* (Fig. 2c). Transmission electron microscopy showed that strains missing *pilA3* and *pilW* or *rntA* and *rntB* are still piliated (Extended Data Fig. 2).

**Fig. 2.**
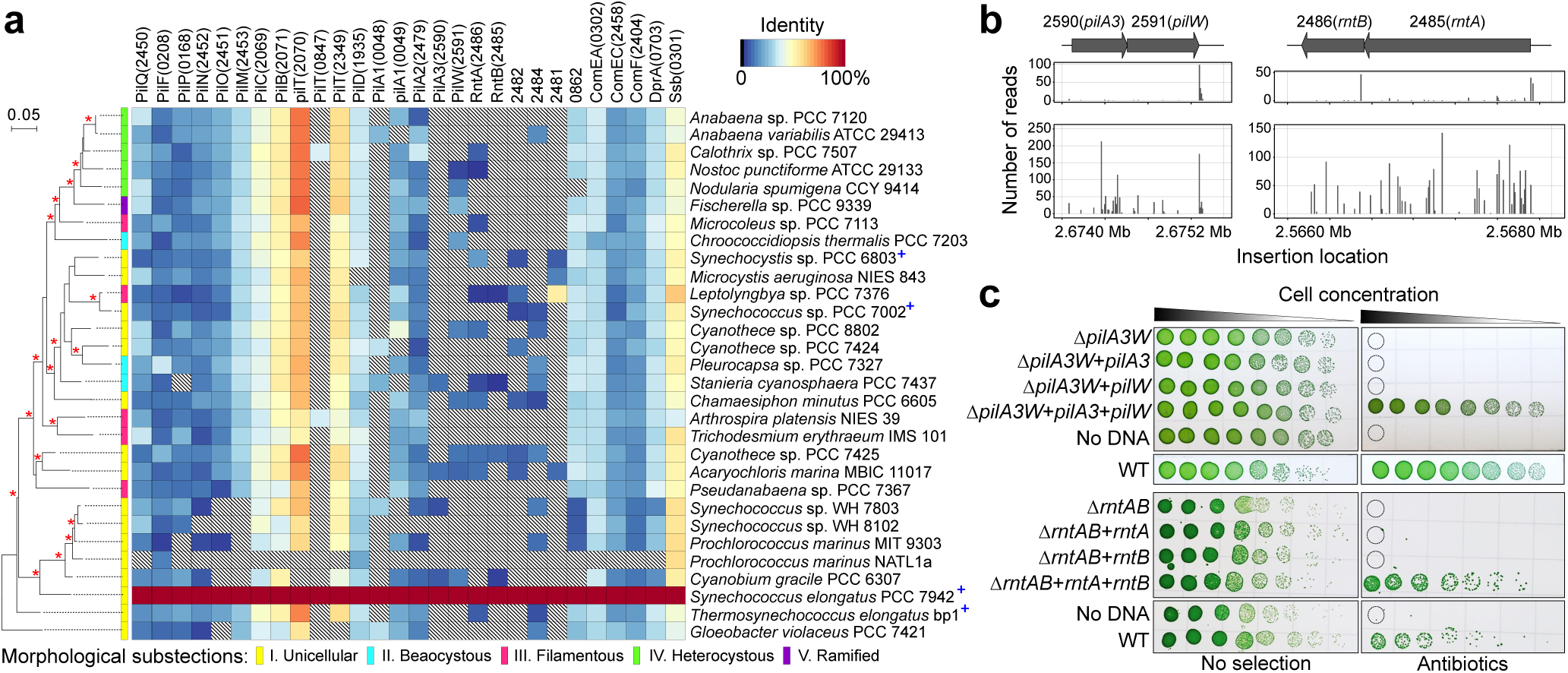
The *S. elongatus* T4PM contains secondary pilins required for natural competence. **a**, Heatmap of protein sequence homologies between T4PM and competence proteins of *S. elongatus* and other strains of cyanobacteria. Hatched cells indicate no homolog in the corresponding strain. Strains organized according to 16S rRNA gene phylogeny, inferred by Maximum Likelihood. Nodes supported by a bootstrap ≥ 70% (n = 500) are marked with a red “*” and strains that are known to be naturally competent are marked with a blue “^+^”. **b**, Genomic organization and RB-TnSeq insertion loci for 2 sets of genes essential for competence. For each these coding regions and flanking sequences, the number of sequencing reads for insertion-mutant barcodes in selective (top boxes) and non-selective control (bottom boxes) conditions is shown for one representative experiment. **c**, Transformation assays performed on deletion and complemented strains of the *pilA3 and pilW*, and *rntA* and *rntB* loci. For each locus, both open reading frames were deleted then complemented separately and together. WT *S. elongatus* PCC 7942 served as a positive control and the fully complemented strains to which no DNA was added served as negative controls. The assays were performed with 3 independent clones for each strain and yielded identical results.

## Dusk and dark

Competence is regulated and transient for most bacteria that are naturally transformable^5^, but no strict competence regulation has been described for *S. elongatus*; however, transformation efficiency increases with the duration of exposure to eDNA and is enhanced in darkness^12^. We hypothesized that natural competence is regulated by the circadian clock, which globally controls gene expression and metabolism in this organism^15,21^.

The RB-TnSeq library screen did not reveal clock genes among the highest determinants of natural competence. The RB-TnSeq data for clock histidine kinase genes *sasA* and *cikA* had mild negative fitness values, but knockout mutants for either of these genes are still naturally transformable^28^. To delve further, we analyzed available circadian and light/dark transcriptomes for *S. elongatus*^19,29^ (Supplementary Table 1). The data clearly indicate that most of the T4PM is circadian controlled and expressed in the morning. In contrast, a subset of genes required for natural competence, including *pilA3, pilW, rntA*, and *rntB*, are circadian with highest expression levels at dusk (Fig. 3a) and are induced by darkness^19,29^ (Fig. 3b).

Electron microscopy of cultures grown in a 12-h light/12-h dark cycle (LD 12:12) confirmed that pili biogenesis occurs largely in the morning (Fig. 3c). Denuded cells incubated from Zeitgeber Time (ZT), the time elapsed since the lights were turned ON, of ZT 0 (dawn) to ZT 5 (late morning) display many pili at the cell surface, while denuded cells incubated during the second half of the night from ZT 18 to 23 (before dawn) remained bald, and cells incubated at the day-night transition from ZT 9 to 14 carried only a few pili. To experimentally test whether natural competence is circadian and dark-induced, quantitative transformation assays on circadian-entrained (LD 12:12) cultures of *S. elongatus* were performed on the second day following their release into constant light (LL). Circadian time (CT) is the biological internal time of entrained cells kept in constant light with CT 0 and CT 12 being subjective dawn and dusk, respectively. For each CT sample, the cells were incubated with eDNA either in the light or in the dark (Fig. 3d). Transformation efficiency peaked at dusk and remained high through the middle of the subjective night. From CT 20 to CT 5, incubation with eDNA yielded no or few transformants. Strikingly, peak transformation efficiency begins at CT 11: the circadian peak for specific essential competence genes, the time when cells anticipate a transition from day to night, and also when darkness starts to strongly impact transformation efficiency.

**Fig. 3.**
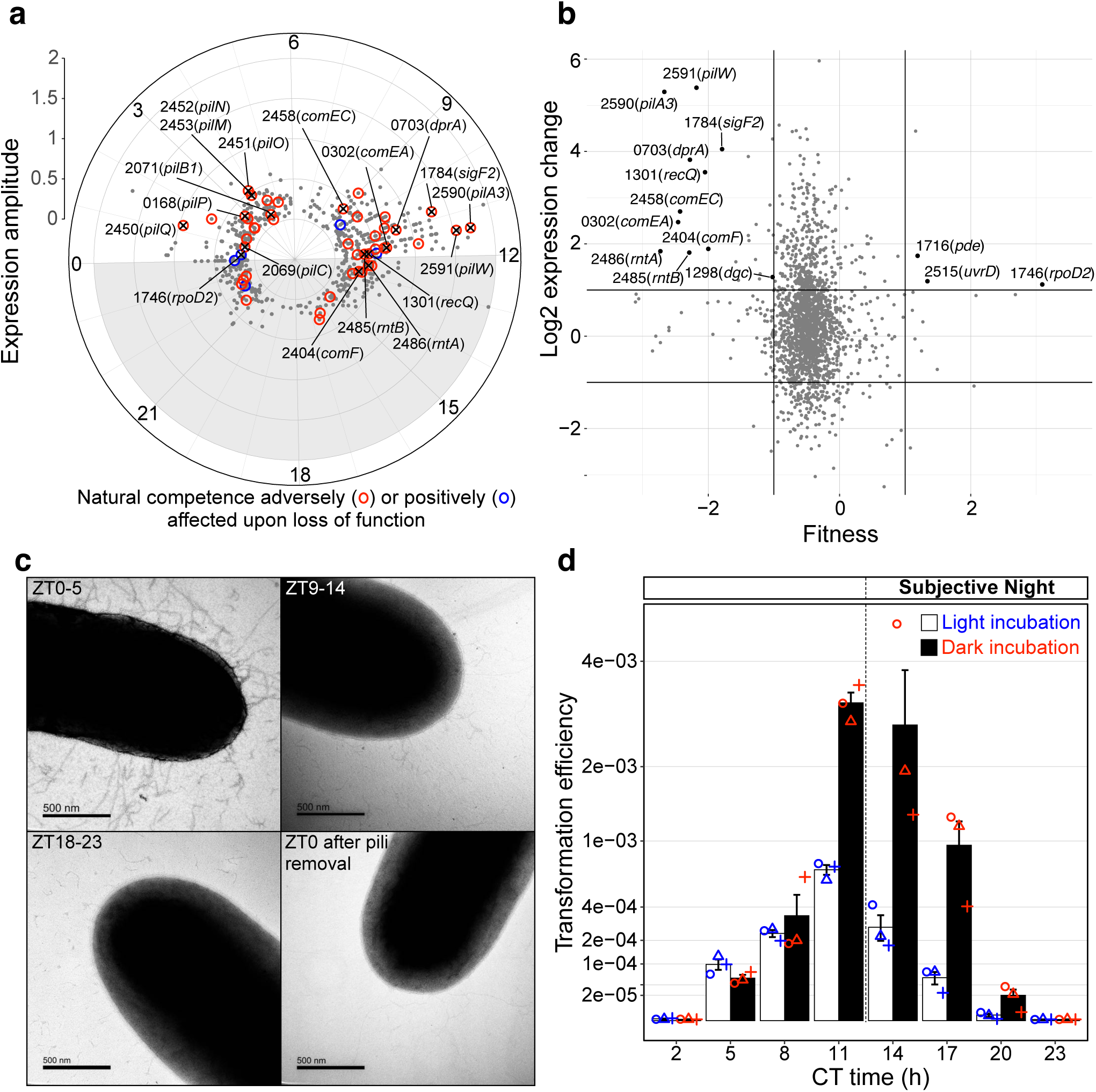
Natural competence is under circadian control and induced in the dark. **a**, Amplitude of circadian-expressed genes reported at peak time^19^. Most of the T4PM genes (including *pilB, M, N, O, P, Q*) are circadian controlled and expressed in the morning while a subset of genes needed for natural competence such as *pilA3, pilW*, and *rntA and rntB*, and the competence genes including *comEA, comEC, comF*, and *dprA*, are circadian with highest expression levels at dusk. **b**, Expression change for genes after a 60-min shade pulse given 8 h after the onset of the day^29^, plotted according to RB-TnSeq fitness values as described for Fig. 1. **c**, Transmission electron micrographs of cells grown in LD and incubated at different time points (Zeitgeber time, ZT time) for 6 hours after removal of their pili. **d**, Transformation efficiency over a 24-h circadian cycle. Entrained cells of WT *S. elongatus* released in LL were incubated with eDNA at different circadian time points (CT time). Efficiencies were calculated as the number of antibiotic-resistant colonies per colony forming unit (CFU) without selection and plotted on a square-root scale to compress large values. Error bars represent standard error from 3 biological replicates (circles, triangles, and plus signs). A similar experiment using a circadian reporter strain of *S. elongatus* and incubated with eDNA carrying a different antibiotic-resistance gene that integrates at a different neutral site yielded similar results (Extended Data Fig. 3).

We predicted that clock-mediated expression of certain dusk-peaking, dark-induced genes is central for the regulation of competence. To investigate the role of the circadian oscillator, we performed quantitative transformation assays at key CT points with two mutants that express phosphomimetic alleles of *kaiC* that lock the oscillator in distinct time-of-day-specific phosphorylation states: a dephosphorylation state more abundant before dawn (KaiC-ET) and a phosphorylation state more abundant at dusk (KaiC-SE)^30^. The expression of *kaiC-ET* (dawn) as the only *kaiC* allele resulted in a strain that lost competence entirely. The *kaiC-SE* (dusk) strain had transformation levels that increased by orders of magnitude when incubated in the dark (∼25 to 500x), achieving high levels of competence at all time points (Fig. 4a). This finding is consistent with induction of competence accompanying the circadian physiology of dusk and early night (Fig. 3d). These results confirm that KaiC orchestrates the timing of natural competence and indicate that the dusk phosphorylation state of KaiC potentiates competence.

**Fig. 4.**
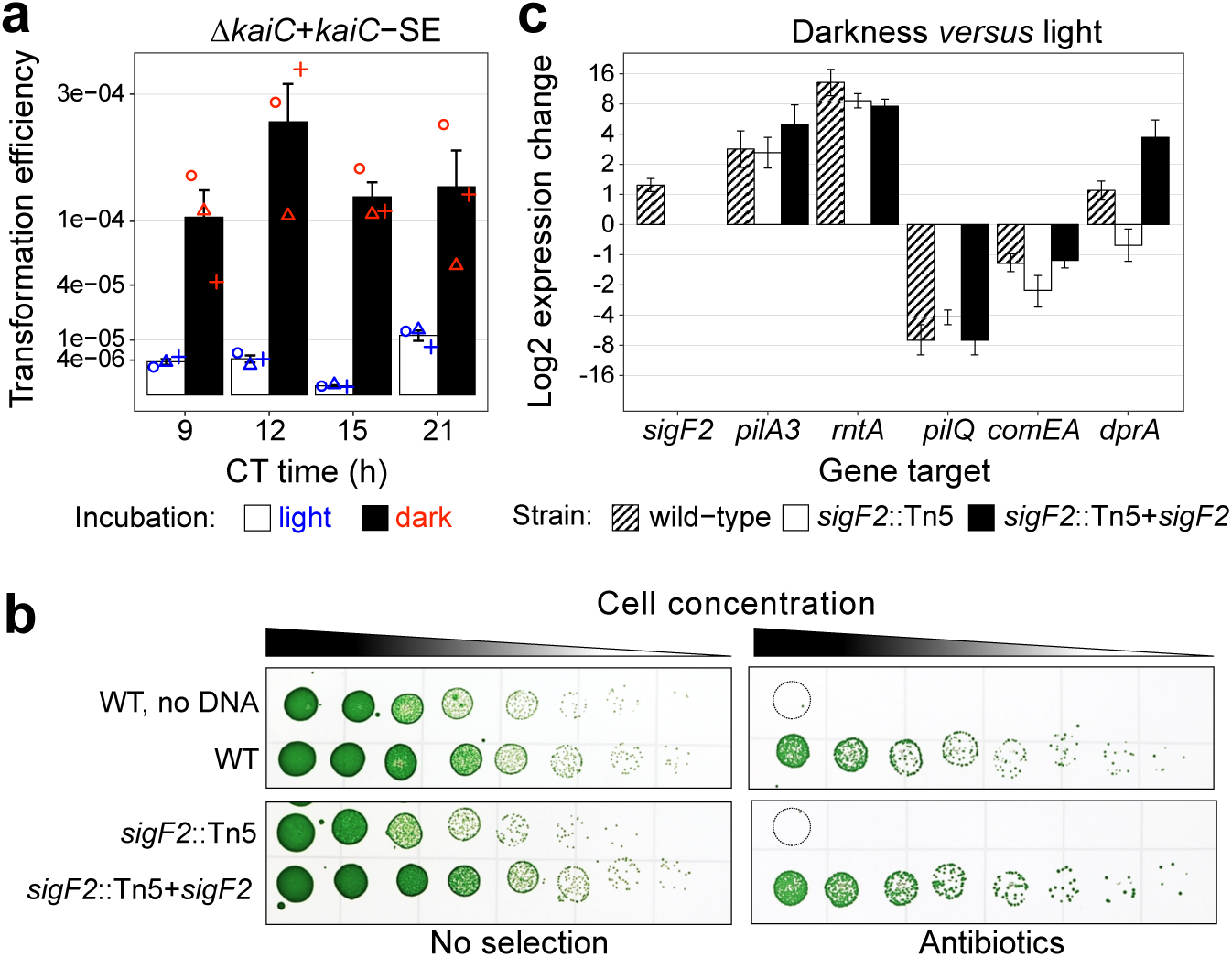
The timing of natural competence is controlled by the phosphorylation state of KaiC and requires SigF2-dependent expression of DprA. **a**, Transformation efficiency at 4 time points over a 24-h circadian cycle of Δ*kaiC* complemented with a mutant *kaiC-SE* phosphomimetic allele that mimics the dusk phosphorylation state of KaiC. Efficiencies were calculated as the number of antibiotic-resistant colonies per CFU without selection and plotted on a square root scale. Error bars represent standard error from 3 biological replicates, with values shown as circles, triangles, and plus signs. **b**, Transformation assays performed on deletion and complemented strains of *sigF2*. WT *S. elongatus* PCC 7942 with and without added eDNA served as controls. The assay was performed using 3 independent clones for each strain, which yielded similar results. **c**, Expression analysis of selected T4PM and competence genes in a *sigF2*-null strain, a *sigF2* complemented strain, and a WT control measured by RT-qPCR on cultures collected two hours after circadian dusk with cells that went into the dark at ZT 12 or were maintained in light. Fold-changes were calculated as 2^-ΔΔCt^ in dark relative to light conditions. Error bars represent standard deviation from 3 biological replicates.

While the induction of dusk genes, including competence genes, is under the control of phosphorylated RpaA (Supplementary Table 1), the mechanism behind increased natural competence in response to darkness is less clear. The expression of hundreds of circadian-regulated genes is affected by changes in light intensity, implicating a network of transcription regulators that is still not well understood^29^. The RB-TnSeq screen revealed the antagonistic importance of two sigma factors in natural competence: *sigF2* and *rpoD2* (Fig. 1a). The role of RpoD2 appears complex and not strictly tied to natural competence^31,32^. SigF2, however, is required for natural competence (Fig. 1a, 4b), has circadian dusk-peaking expression that increases with darkness (Fig. 3a,b), and is a target for RpaA and RpaB^19,29^, which are both implicated in the regulation of circadian genes by light availability. SigF2 is paralogous to SigF, which in other cyanobacteria is associated with pili formation and motility^33^. Although, SigF is also tied to the T4PM in *S. elongatus* based on previous RB-TnSeq screens^34^, those screens did not reveal a role for SigF2. Thus, we asked whether SigF2 contributes to the dark induction of natural competence. The relative transcript levels of selected genes required for competence, and (except *pilQ*) that have been reported to be dark-induced were investigated by RT-qPCR for the wild type (WT), a *sigF2* knockout, and a complemented *sigF2* strain (Fig. 4c). To determine whether key genes are induced by darkness, LD-entrained cells were either maintained in the light or transferred to darkness at ZT 12 and sampled for RNA 2 h later, which corresponds to the peak of natural competence in LD. For the WT, our results sampled 2 h after dusk generally agreed with data sampled 3-4 h before dusk^29^ that *sigF2, pilA3, rntA*, and *dprA* are dark induced. However, *comEA* expression decreased in the dark after dusk, whereas it increased in the dark before dusk^29^, suggesting that *comEA* dark induction is tightly controlled by the clock. Notably, *dprA* induction did not occur in the absence of SigF2, consistent with evidence for *sigF2*-dependent expression of *dprA* in a recent transcriptomics dataset obtained in LL^20^. The complementation of *sigF2* from a constitutive promoter restored *dprA* dark induction and resulted in overexpression of *dprA.* The downregulation of PilQ at dusk in darkness supports the idea that fewer T4PM are produced in early night, while specific competence-related T4P are made at this time. The results suggest that *sigF2*, which responds to light availability^29^, participates in a complex regulatory network that mediates dark-induced competence at dusk, and implicates *dprA* as an important target (Fig. 4d).

## Seasonal photoperiod

The regulation of competence by the clock and darkness is reminiscent of a model of external coincidence proposed by Erwin Bunning in 1936^35^. This model explains how the underlying circadian cycle adjusts to different seasonal photoperiods to change the abundance at dusk or dawn of components whose activity is directly or indirectly affected by the presence of light. For example, external coincidence underlies the timing of daily hypocotyl elongation in *Arabidopsis*^36^. Although cyanobacterial growth varies over seasons in the environment^37^, no studies address a contribution of photoperiodic regulation or a role for the internal circadian clock. We asked whether the daily pattern of natural competence is affected by changes in photoperiod that mark seasonal variations.

Levels of competence were measured every 2 h over a 24-h time course in LL and LD using cultures of *S. elongatus* entrained with distinct photoperiods: LD 12:12 for regular days and 16:8 for long days associated with summer; cells did not thrive in an LD 8:16 winter photoperiod. To maximize transformation levels in LL, incubations with eDNA were performed in darkness; while in LD, incubations were performed in either light or dark according to the diel condition at the time of transformation. The results of this experiment (Fig. 5) agreed with those presented in Fig. 3d and with the prior finding that the clock tracks midday^38^. In LL (Fig. 5a,c), the peak and trough times of competence occur 2 h later in long days versus regular days, whereas the onset of night occurs 4 h later. Thus, peak transformation efficiency occurs at the onset of subjective night in regular days and 2 h prior to subjective night in long days. By extrapolating these data over a few cycles, the mfourfit method^39^ also calculated a shift of 2.1 h between regular and long days. In regular days the amplitude of transformation efficiency was greater than in long days, falling off sharply on either side of the peak, whereas a broad peak of competence in long days resulted in high competentence during a time window of 8 h versus only 4 h in regular days. In LD (Fig. 5b,d), a dramatic difference in competence occurs between day and night. In regular days, competence peaked to almost 5% of cells being transformed during the night but was almost entirely suppressed from dawn until dusk. In long days, a small fraction of the cells remains competent at dawn. The peak of competence occurs 2 to 4 h after nightfall in both regular and long days. In long days, the peak time is delayed by 4 h compared to regular days. To further illustrate the cooperative effect of darkness on natural competence, additional assays were performed on cultures maintained for 4 h in the light after anticipated nightfall and in the dark at the onset of the day (Fig. 5b,d, white bars below or next to black bars). In both photoperiods, natural competence remains low after anticipated nightfall if the cells are maintained in light conditions, demonstrating the need for dark induction; furthermore, as in LL, darkness has little effect on natural competence if not aligned with the circadian peak.

**Fig. 5.**
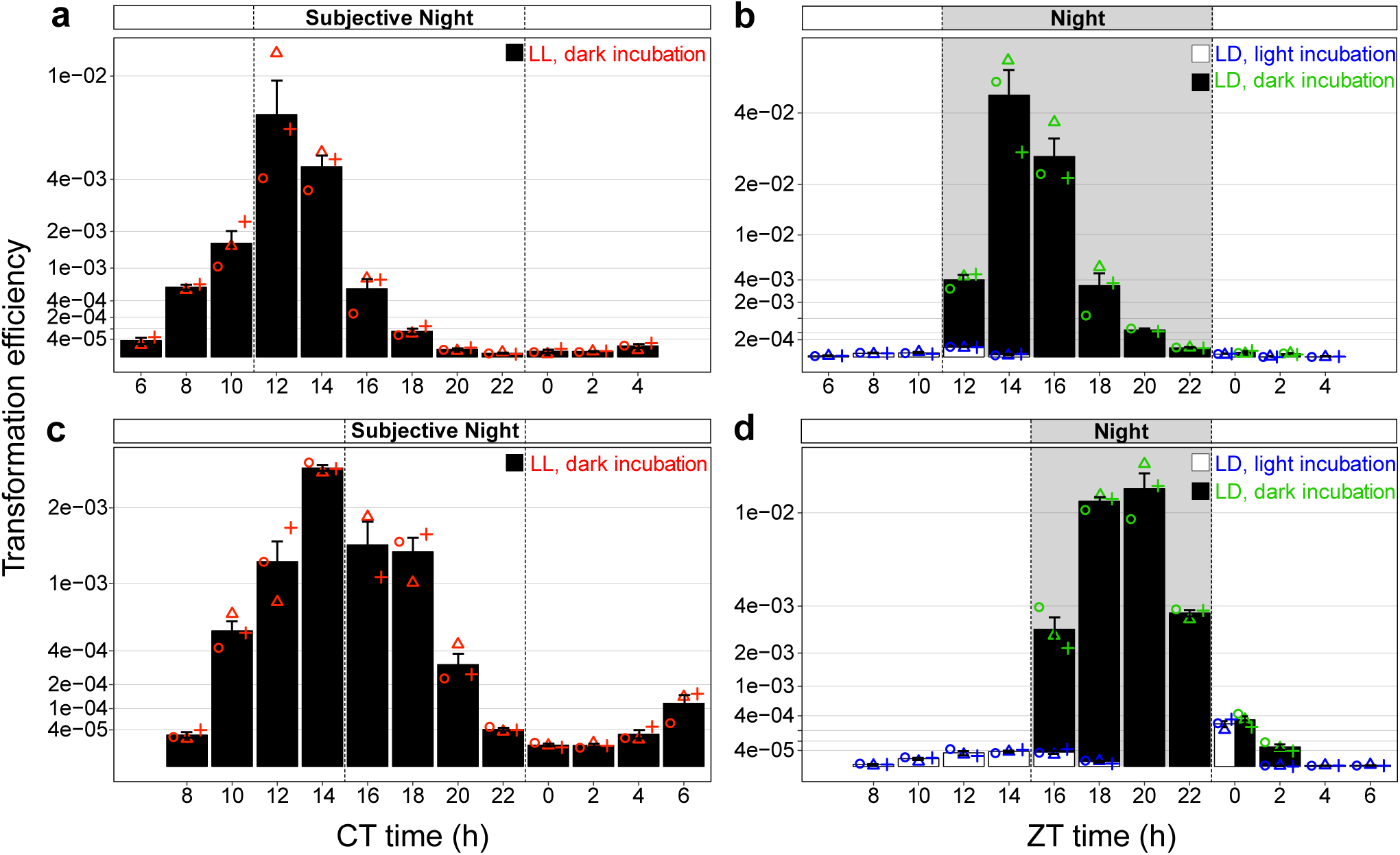
Natural competence is under circadian control, requires external coincidence with darkness, and responds to changes in day length. Transformation efficiency over a 24-h circadian cycle of cultures entrained with **a, b**, 12-h day length, and **c, d**, 16-h day length under **a, c**, constant light, and **b, d**, light-dark conditions. Efficiencies were calculated as the number of antibiotic resistant colonies per CFU without selection and plotted on a square-root scale. Error bars represent standard error from 3 biological replicates, with values shown as circles, triangles, and plus signs. White and black bars are either overlaid or shown side-by-side to make both values visible.

## Discussion

Cyanobacteria are important photosynthetic primary producers in many environments^14^ and have great potential as platforms for the production of renewable biochemicals^8^, so understanding their genetic exchange mechanisms is of ecological and practical importance. In-depth studies of natural competence have largely focused on bacterial pathogens^*3,40-42*^, so little is known about the process in other prokaryotic phyla. In cyanobacteria, some of the conserved T4PM and competence proteins had been shown to be essential for competence^10,43^. Our work experimentally extends this knowledge to encompass all proteins that are required for the process including those that could not be identified solely by bioinformatics^11^. Beyond conserved T4PM and competence proteins, we also identified hypothetical proteins and PilA3, a rare minor pilin, which are required for natural transformation of *S. elongatus*. Similarly, minor pilins were found to be necessary for floc formation in the cyanobacterium *Synechocystis* sp. PCC 6803^44^.

Although a fundamental blueprint for DNA uptake and processing may be conserved among bacteria, the regulatory mechanisms and inducing cues vary among phyla^5,6^. The finding of a daily rhythm of natural competence, controlled by the circadian clock and nightfall, makes physiological sense for the phototrophic cyanobacteria, invoking recurrent general themes of nutritional status and stressful conditions^5^ as drivers of competence. Their internal clock allows cyanobacteria to anticipate the daily change in light and dark, which is important as they rely upon light to drive photosynthetic metabolism and growth, and must adapt their physiology daily to darkness^45^. While the circadian rhythmicity of competence can be tied directly to the phosphorylation state of KaiC and the RpaA-dependent expression of dusk-peaking genes^19,21^ that are essential for natural competence, the network of transcriptional regulators beyond RpaA and RpaB^29^ is unmapped. We found that the alternative sigma factor SigF2, which is a target of RpaA and RpaB^19,29^, factors into the dark-inducing mechanism, likely through its role in the expression of DprA, which loads ssDNA on RecA prior to recombination^3,5^.

A convergence of environmental signals and internal circadian rhythms is a hallmark of photoperiodic flowering in plants^46^ and seasonal reproductive rhythms in mammals^47^, and controls daily growth rhythms in *Arabidopsis*^36^. Similarly, the circadian peaking of gene expression required for natural competence is synergistic with darkness in rendering cyanobacterial cells competent. The circadian clock controls numerous processes in *S. elongatus*^7^ including cell division. According to the prevailing model, the clock selectively inhibits cell division during a temporal window during the night, and cells divide freely at other times of the day^23,24^. A recent study further showed that instead of solely applying an open-close gate on division, cells must integrate internal and external cues to decide when to divide^48^. The evolutionary pressure for circadian gating of cell division or competence is not obvious, but they may be related, as the timing of maximal natural competence roughly corresponds to the window during which cell division is disallowed. Growth arrest coregulated with competence has been observed in other model organisms^6^. Moreover, the *S. elongatus* chromosome undergoes a clock-dependent compaction cycle, with chromosomes maximally compacted around dusk and decompacting during the night^22^. These rhythms suggest that there may be a limited time during the day when eDNA taken up from the environment could be productively incorporated into the genome by homologous recombination. Why did it take so long to discover the circadian rhythm of competence in *S. elongatus*, which is famous for its robust clock? The strain is so compliant as a model organism that daily variations were not notable until queried rigorously^12^. For convenience cultures are grown in LL, where they remain rhythmic, but the population exhibits variations in clock phasing such that competent cells are always present during routine laboratory growth, and the transformation of such cultures always leads to more transformants than are required for most genetic experiments.

## Methods

### Strains, plasmids, and molecular methods

*Escherichia coli* strains were grown at 37 °C in lysogeny broth (LB, Lennox) liquid culture or on agar plates, supplemented as needed with: 100 µg ml^-1^ ampicillin (Ap), 20 μg ml^-1^ spectinomycin (Sp) plus 20 μg ml^-1^ streptomycin (Sm), 50 μg ml^-1^ spectinomycin (Sp), 15 μg ml^-1^ gentamycin (Gm), 17 µg ml^-1^ chloramphenicol (Cm), 50 µg ml^-1^ kanamycin (Km), and 50 µg ml^-1^ nourseothricin (Nt). Unless noted otherwise *S. elongatus* PCC 7942 and its derivative strains were grown in BG-11 medium^49^ as liquid cultures with continuous shaking (125 r.p.m.) or on agar plates (40 mL, 1.5% agarose) at 30 °C under continuous illumination of 200 μmol photons m^-2^ s^-1^ from fluorescent cool white bulbs. Culture media for recombinant cyanobacterial strains were supplemented as needed with 2 μg ml^-1^ Sp plus 2 μg ml^-1^ Sm, 2 μg ml^-1^ Gm, 7.5 µg ml^-1^ Cm, 5 µg ml^-1^ Km, and 5 µg ml^-1^ Nt.

Recombinant strains of *S. elongatus* were constructed by natural transformation using standard protocols^27^. Briefly, liquid cultures were grown to an OD_750_ of 0.5-0.6. Cells were pelleted by centrifugation at 4,500*g*, washed once with BG-11 medium and once with 10 mM NaCl, then resuspended in BG-11 to a concentration of 1 × 10^9^ cells/ml. Samples (200 µl) were incubated in darkness for 16 h with 500-1000 ng of plasmid DNA. Transformation reactions were then plated on BG-11 agar with appropriate antibiotics and incubated for 4 to 5 days under continuous illumination until isolated colonies appeared. Complementation and overexpression strains were constructed by expressing a gene(s) ectopically in *S. elongatus* chromosomal neutral sites: NS1, NS2, and NS3^27,50^. To complement genes essential for natural transformation, such as *pilA3* and *pilW, rntA* and *rntB*, or *sigF2*, we first constructed strains with one or both genes expressed ectopically and then deleted the native loci. Complete segregation of the deleted loci was PCR-verified, and 3 independent clones were picked as biological replicates for subsequent experiments. Plasmids were constructed using the GeneArt Seamless Cloning and Assembly Kit (Life Technologies) and propagated in *E. coli* DH5α or TOP10 with appropriate antibiotics. Gene knockouts in *S. elongatus* were constructed using *S. elongatus* UGS library plasmids^51^ or with plasmids engineered with CYANO-VECTOR assembly^52^. Strains and plasmids used in this study are listed in Supplementary Table 2.

### RB-TnSeq library screen for natural competence genes

To screen the *S. elongatus* RB-TnSeq library^25^ for genes involved in natural competence, an aliquot of the library archived at -80 °C was thawed quickly and regrown as previously described^25^ to an OD_750_ of 0.3. A sample of the library prior to transformation (T_0_) was collected to determine the population baseline. Then natural transformations were performed as described above except that cells were incubated with 1 µg of exogenous plasmid DNA in low light (10 μmol photons m^-2^ s^-1^) instead of complete darkness to preserve mutants that do not survive light-dark cycles. For each experiment, 10 transformation reactions were prepared and plated on selective (Nt or Sp+Sm) and non-selective (control condition) BG-11 agar plates and incubated for 4 to 5 days under continuous illumination until isolated colonies appeared. Three experiments were performed using plasmid DNA carrying either an Sp/Sm resistance gene (pAM5329) that recombines into the *S. elongatus* chromosome at NS1 (performed twice) or an Nt resistance gene (pAM5544) that recombines at NS2^27^. For each experiment, 70,000 to 250,000 colonies were collected, pooled, and stored at -80 °C for genomic DNA extraction and barcode sequencing^25^.

Sequencing reads of the barcodes were curated to keep only barcodes located within the middle 80% of each coding sequence; genes not represented by at least three bar codes in different positions or without at least 15 T_0_ reads across replicates were removed from the analysis^26,53^. This curation resulted in 82,495 barcodes (also named strains) for the experimental conditions, 90,997 barcodes for the controls, and 99,872 barcodes for the T_0_ samples distributed across 1,885 genes. For each gene, a fitness value that describes how the loss of function affects natural competence and a corresponding statistic were estimated in R as described previously^26,53^.

### Construction of the heatmap of protein sequence homologies and the 16S rRNA gene phylogeny

Proteins that belong to T4PM including secondary pilins, that carry a pilin-like signal peptide, or are recognized as essential for natural competence in other bacteria were initially identified in *S. elongatus* based on its genome annotation, the available literature^11,54^, or sequence homologies using proteins with known functions from other organisms. For each protein encoded by *S. elongatus*, BLASTp searches were conducted against selected cyanobacterial strains using BioBIKE (http://biobike.csbc.vcu.edu/)^55^ to identify the best reciprocal hit with an e-value greater than or equal to 0.001. The heatmap was then generated based on the sequence identities using the heatmap.plus R package.

A multiple alignment of 16S rRNA gene sequences and the phylogenetic tree were constructed using the software package MEGA. The alignment was generated with MUSCLE^56^ implemented in MEGA using default parameters. Positions that can be used reliably in a phylogenetic analysis were extracted with Gblocks^57^ at settings that allowed the most relaxed selection of blocks and covered 1,338 positions of the alignment. Evolutionary history was inferred by using the Maximum Likelihood method based on the Kimura 2-parameter model^58^ implemented in MEGA. The tree with the highest log likelihood (−9,284.19) is shown. The percentage of trees in which the associated taxa clustered together was calculated for each node and indicated with an “*” when above 70%. The tree is drawn to scale, with branch lengths measured in the number of substitutions per site.

### Transformation assays of non-synchronized cells

Knockout, complementation, and overexpression strains for selected genes identified as important for natural competence, as well as appropriate control strains, were grown as described earlier and subjected to semi-quantitative transformation assays. Transformation reactions (cells incubated with exogenous DNA, eDNA) were prepared as described above using the subject strain and appropriate plasmid DNA (pAM5328, pAM5544, or pAM5554) that carries an antibiotic resistance gene, and that will recombine into a neutral site. For each strain, assays were performed with 2 or 3 independent clones. Transformation reactions were serially diluted and spotted (8 spots of 4.5 µl) on agar plates (10 cm square petri dishes) without antibiotic to determine total colony forming units (CFU), and with antibiotic to select for transformed clones.

### Transformation assays of light/dark entrained cells

To obtain quantitative transformation efficiency levels over a circadian time course, our transformation protocol was modified to account for the time sensitivity of these experiments and the effect of light vs dark incubation with eDNA.

For the experiments presented in Fig. 3d and Extended Data Fig. 3, pre-cultures of *S. elongatus* entrained in 12-h light/12-h dark cycles (LD 12:12) were used to inoculate triplicate 100-ml cultures to an OD_750_ of 0.1 in BG-11 medium at the onset of light. They were grown using stirred flasks for 2 days in LD 12:12 and then transferred to constant light (LL) on the third day. On the second day in LL, quantitative transformation assays were performed every 3 h using 200 µl of cells, washed once with BG-11, concentrated to 1 × 10^9^ cells/ml and incubated for 3 h with 500 ng of plasmid DNA either in the light or in the dark. The incubations in the dark were limited to 2 h followed by 1 h in the light to reduce the chance of resetting circadian phase. Transformation reactions were stopped by serially diluting and inoculating samples (25 µl) onto selective or non-selective BG-11 in 12-well plates (3 ml/well, 1.5% agar). The plates were incubated for 5 days at 30 °C in LL until colonies had appeared. For each plate, wells with isolated colonies were photographed (typically 2 wells for the selective medium and 1 well for the control). Pictures were processed with ImageJ and curated manually to produce binary images without background and where adjacent colonies were separated. The number of colonies was determined with ImageJ. Transformation efficiencies were calculated as the number of antibiotic-resistant colonies per CFU without selection. This experiment was first performed with WT *S. elongatus* using pAM5544, which carries a Nt resistance gene and integrates at NS2 by homologous recombination (Fig. 3d). This experiment was later repeated with a circadian reporter strain, AMC1300, using pAM5328, which carries a Gm resistance gene and recombines at NS3 (Extended Data Fig. 3a).

For the experiments presented in Fig. 4, with the exception of AMC2109, which is dark sensitive and was maintained in LL, precultures grown in LD 12:12 were used to inoculate 100-ml cultures to an OD_750_ of 0.1 in 250-ml Falcon tissue culture flasks bubbled with air (0.1 L/min). Cultures were grown in LD 12:12 for 3 days until they reached an OD_750_ of ∼0.5. On the third day at nightfall, cells were concentrated to 5 × 10^8^ cells/ml and distributed as 200-µl aliquots in a white 96-well plate with a clear bottom. The plate was placed in the incubator between 2 light sources to minimize shading effects at the bottom of the wells and to maintain a light intensity of 200 µmol photons m^-2^ s^-1^ on both sides of the plate. At that time the light sources were turned OFF until dawn, 12 h later. Afterwards, the 96-well plates were maintained in LL and quantitative transformation assays were performed at 4 time points. For each time point, 200 ng of plasmid DNA was added to the wells, the cells and eDNA were mixed by slowly pipetting up and down, and samples were incubated in the dark for one hour by covering the top and bottom of the wells with opaque aluminum sealing tape. After incubation, transformation reactions were serially diluted, inoculated into 12-well plates, incubated for several days, photographed, and processed to determine colony counts and transformation efficiencies as described above.

For the experiments on day length presented in Fig. 5, pre-cultures of *S. elongatus* grown in LD 12:12 and LD 16:8 were used to inoculate 100-ml cultures to an OD_750_ of 0.1 (LD 12:12) or 0.05 (LD 16:8), which were grown in tissue culture flasks as described above. On the third day at nightfall, the cells were concentrated and distributed in 96-well plates, which were incubated in LD 12:12 or LD 16:8 starting with a dark period of 12 and 8 h for the cells entrained in regular or long days, respectively. Quantitative transformation assays were performed every 2 h, as described for the previous experiment. The time course started at the middle of the day, at ZT 6 (6 h after dawn for cultures entrained in regular days) and ZT 8 for the cultures entrained in long days. Because the incubators were set to constant light, dark conditions were achieved by covering the bottom and top of wells with opaque aluminum sealing tape for 12 h (regular days) or 8 h (long days). Transformation reactions were incubated for one hour either in the dark or in the light. After incubation, the transformation reactions were serially diluted, inoculated into 12-well plates, and incubated as described above. Triplicate plates were stacked and rotated every day to equalize the total amount of light that each plate received. Wells were photographed and the pictures were processed to determine colony counts as described above. An average of 73 colonies/well were counted for 466 wells. For this experiment, 56 transformations were performed in triplicate (168 transformations in total).

### RT-qPCR

Precultures grown in LD 12:12 were used to inoculate 100-ml cultures to an OD_750_ of 0.1 in 250 ml Falcon tissue culture flasks bubbled with air (0.1 L/min). Cultures were grown in LD 12:12 for 3 days until they reached an OD_750_ of ∼0.5. On the 3^rd^ day at nightfall, each culture was divided in 2 subcultures with one maintained in constant light and the second put into the dark, and two hours later the cells were collected, poured over ice in a 50 ml falcon tube and spun down at 4 °C for 10 min at 4,500*g*. Cell pellets were frozen at -80 °C. RNA was extracted from triplicate cultures for each strain and condition. Cell pellets were resuspended in 700 µl of TRIzol and incubated for 5 minutes at 95-100 °C followed by 5 minutes at room temperature. The mixture was spun down (16,000*g*) for 5 min at 4 °C and the supernatant extracted with 700 µl (1 volume) of 100% EtOH at 4 °C. All following steps were performed at 4 °C using RNase-free tubes and tips. RNA was isolated using the Direct-zol Quick RNA miniprep Kit (Zymo Research) following the manufacturer’s instructions. RNA concentration ranged from 900 -960 ng/µl as measured with a Nanodrop (Thermo Fisher Scientific). RNA was treated with DNaseI (RNase free, Thermo Fisher Scientific) following the manufacturer’s instructions and stored at -80 °C. cDNA was synthesized from 500 ng of RNA template per sample with the High-Capacity RNA-to-cDNA Kit (Applied Biosystems). RT-qPCR reactions were performed in triplicate with the Biotool 2x SYBR Green qPCR master mix (Low ROX) reagents following the instructions in The Applied Biosystems StepOne Real-Time PCR Systems kit. Primer sequences and target genes are listed in Supplementary Table 3. Gene expression fold-changes in cultures incubated in the dark were calculated relative to cultures incubated with light (Fig. 4c) using the 2^−ΔΔCt^ method^59^ with *rnpB* (Synpcc7942_R0036) as the reference.

### Electron microscopy

Small aliquots (200 µl) of *S. elongatus* cultures entrained in LD 12:12 were fixed with 0.5% glutaraldehyde for 5-10 min at room temperature. Sample grids (Formvar/Carbon 100 mesh, Copper) were floated for 2 min on a drop of the fixed cells, rinsed 3 times by floating the grids on drops of water, and floated on a drop of 1% uranyl acetate for 2 min to stain the cells. Excess uranyl acetate solution was removed with filter paper and grids were air dried for a few minutes before electron microscopy. EM was performed with a JEOL 1200 EX II TEM equipped with a Gatan Orius 600 7-megapixel bottom-mount digital camera (2.7k x 2.7k).

To determine the timing of pili biogenesis, precultures of *S. elongatus* grown in LD 12:12 were used to inoculate 100-ml cultures to an OD_750_ of 0.1 in 250-ml Falcon tissue culture flasks bubbled with air (0.1 L/min), which were grown in LD 12:12 for 3 days until they reached an OD_750_ of ∼0.5. On the third day at nightfall, cells were concentrated (200 µl at 10^8^ cells/ml) and distributed in a 96-well plate; the plate was placed in the dark incubator until the next morning. At ZT 0, ZT 9, and ZT 18 the concentrated cells were spun down (4500*g*), washed twice with BG-11 to remove their pili, resuspended with 200 µl of BG-11 in the 96-well plate, and returned in the incubator for 6 h until the cells were prepared for electron microscopy.

### Circadian bioluminescence monitoring

*S. elongatus* strains expressing P_*psbA*_-*luxCDE* and P_*kaiBC*_-*luxAB* or P_*furA*_-*luxAB* were grown at 30 °C for 2 or 3 cycles in LD 12:12 in the conditions described for the corresponding transformation assays (Extended Data Fig. 3). At the onset of night on the day before being released into constant light, 20 µl of cells were distributed in 96-well agar plates prepared as described previously^60^. On the day of the transformation assays the plates were placed in a TECAN Infinite 200 Pro equipped with a stacker microplate handling system where plates were maintained at 30 °C in LL. Bioluminescence readings were recorded every 3 h for 48 h. Data were analyzed with BioDare2 using linear detrending and normalized to the mean (https://biodare2.ed.ac.uk/)^39^.

## Data availability

RB-TnSeq, transcriptomics data, descriptions of plasmids and strains and RT-qPCR primers are available as supplementary tables; additional supporting material for the findings of this study are also available in the extended data.

## Acknowledgments

We thank D. Welkie, C. Sancar, and R. Simkovsky for their input on the experimental design and data analyses, B. McKnight, L.C. Lowe, and C. Peterson for assistance with strain construction and transformation assays. We thank T. Meerlo at the UC San Diego electron microscopy facility and K. Jepsen at UC San Diego IGM sequencing facility for technical support. Funding was provided by the National Institute of General Medical Sciences of the National Institutes of Health under award number R35GM118290 (to S.S.G.) and R01GM118815 (to J.W.G). The content is solely the responsibility of the authors and does not necessarily represent the official views of the National Institutes of Health. This material is also based upon work supported by the National Science Foundation under Grant Number IOS-1754894. Any opinions, findings, and conclusions or recommendations expressed in this material are those of the author(s) and do not necessarily reflect the views of the National Science Foundation.

## Author contributions

A.T, J.W.G, and S.S.G. conceived of and designed the project. A.T. led or performed the experimental work and analyzed the data, C.E. performed the RB-TnSeq transformation screens and helped with strain constructions and preliminary transformation assays. B.E.R. provided the RB-TnSeq library and obtained barcode sequences. Y.Y. performed RT-qPCR experiments. S.A.R. contributed to RB-Tn-Seq analytic tool. A.T, J.W.G., and S.S.G. wrote the manuscript, which was reviewed and edited by all authors.

## Competing interest

The authors declare no competing interests.

## Additional information

**Extended data** is available for this paper at

**Supplementary information** is available for this paper at

**Reprints and permissions information** is available at

---

**Extended Data Fig. 1.**
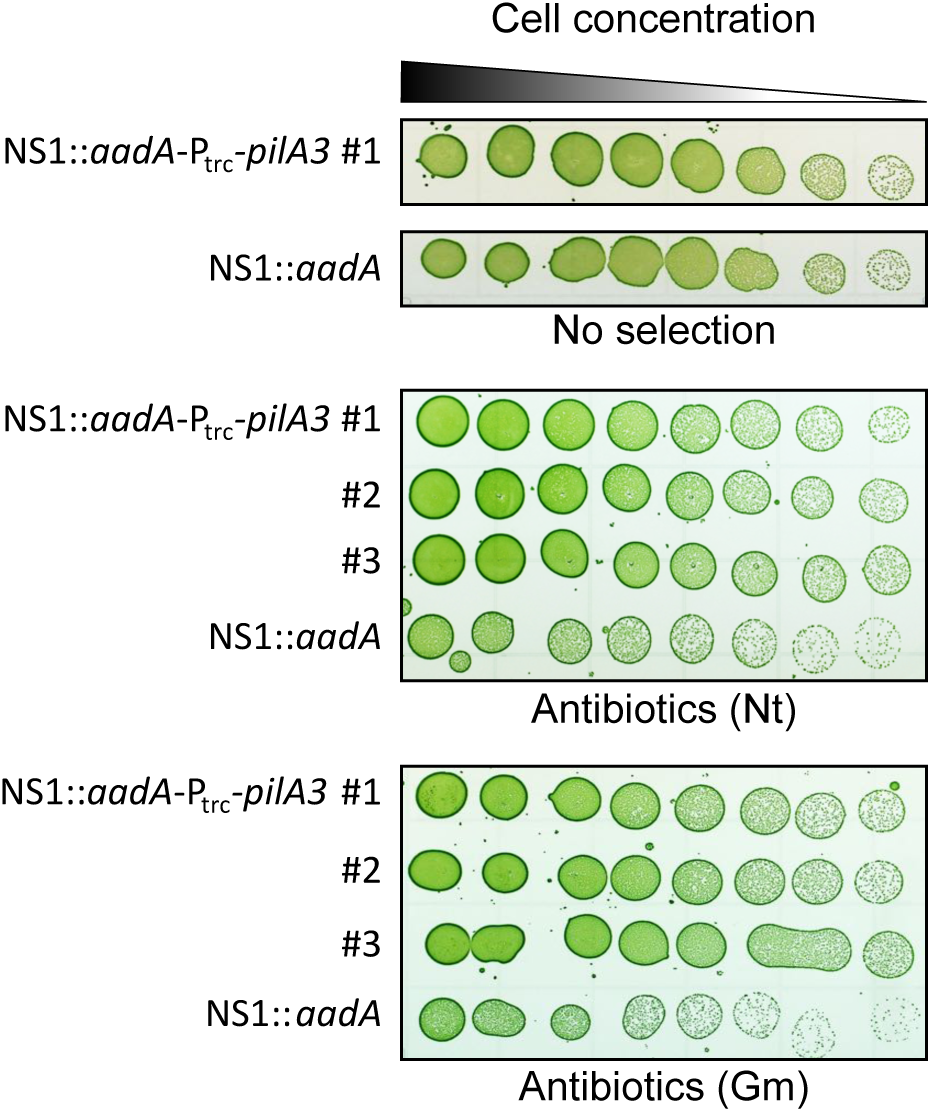
Overexpression of *pilA3* results in increased competence. Semi-quantitative transformation assays performed on a *pilA3* overexpression strain. The assays were performed on 3 independent clones with plasmids that carry different antibiotic resistance genes and target distinct neutral sites (NS) in the *S. elongatus* chromosome.

**Extended Data Fig. 2.**
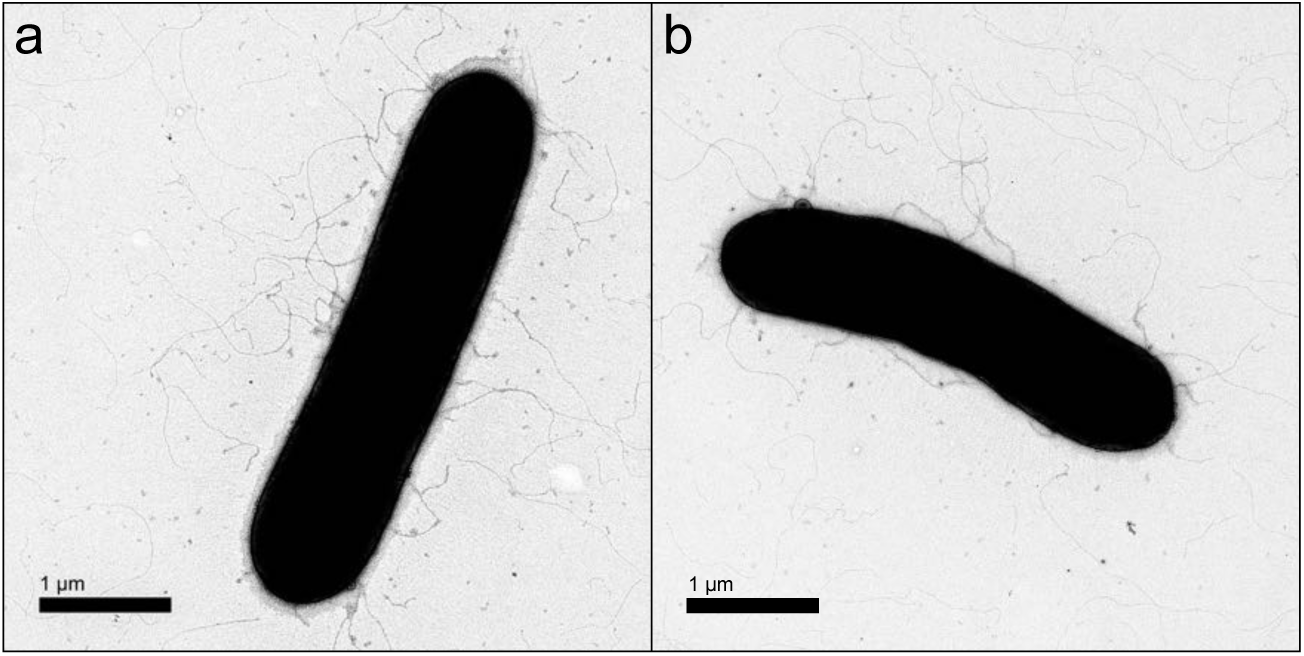
*S. elongatus* remains piliated upon deletion of *pilA3* and *pilW* (AMC2547) or *rntA* and *rntB* (AMC2543). Transmission electron microscopy pictures of **a**, strain AMC2547 and **b**, strain AMC2543.

**Extended Data Fig. 3.**
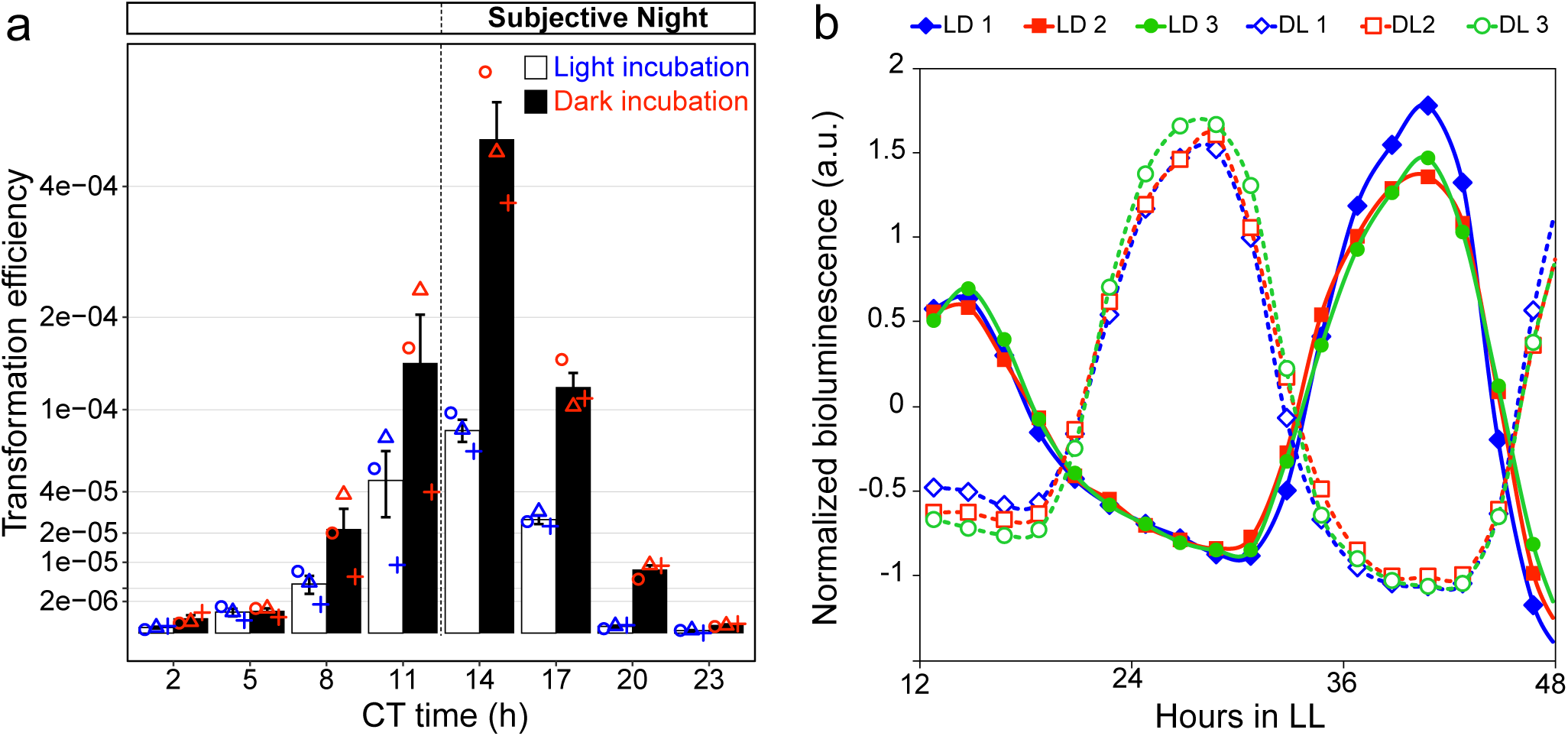
Natural competence is circadian and induced by darkness. **a**, Transformation efficiency in *S. elongatus* circadian reporter strain AMC1300 over a 24-h circadian cycle. **b**, Circadian rhythmicity of gene expression in cultures of *S. elongatus* AMC1300 used for the transformation assays. Bioluminescence from strain AMC1300 is produced by a P_*kaiB*_-*luxAB/P*_*psbA*_*-luxCDE* reporter system.

